# Characterizing molecular flexibility by combining lRMSD measures

**DOI:** 10.1101/379784

**Authors:** F. Cazals, R. Tetley

## Abstract

The root mean square deviation (RMSD) and the least RMSD are two widely used similarity measures in structural bioinformatics. Yet, they stem from global comparisons, possibly obliterating locally conserved motifs. We correct these limitations with the so-called *combined RMSD*, which mixes independent lRMSD measures, each computed with its own rigid motion. The combined RMSD can be used to compare (quaternary) structures based on motifs defined from the sequence (domains, SSE), or to compare structures based on structural motifs yielded by local structural alignment methods.

We illustrate the benefits of combined RMSD over the usual RMSD on three problems, namely (i) the analysis of conformational changes based on combined RMSD of rigid structural motifs (case study: a class II fusion protein), (ii) the calculation of structural phylogenies (case study: class II fusion proteins), and (iii) the assignment of quaternary structures for hemoglobin. Using these, we argue that the combined RMSD is a tool a choice to perform positive and negative discrimination of degree of freedom, with applications to the design of move sets and collective coordinates.

Combined RMSD are available within the Structural Bioinformatics Library (http://sbl.inria.fr).

## 1 Introduction

### 1.1 RMSD, lRMSD and their variants

The problem of geometrically comparing two point sets of the same cardinality, assuming a one-to-one correspondence between the points, has long been recognized as central in science and engineering. It is in general desired to perform a comparison oblivious to rigid motions, which prompts a solution computing concomitantly the geometric similarity measure and the associated optimal rigid motion. The most celebrated solution to this problem is the so-called least root mean square deviation (lRMSD)[1], namely the RMSD of positions upon applying the optimal rigid motion. This number, which is usually expressed in Å. (We note that the lRMSD is a coordinate RMSD, not the be confused with the *RMSD_d_*, namely the RMSD of internal distances.) In the sequel, we review previous work, by restricting ourselves to structural bioinformatics.

Several strategies to compute the lRMSD [1, 2, 3] or its weighted variant [4, 5] were developed long ago, and it was also noted that the lRMSD induces a metric [6]. Owing to these properties, the lRMSD has been one of the most used similarity criteria in structural biology and bioinformatics. On the other hand, several limitations prompted developments from the design and computational perspectives.

On the design side, efforts were made to circumvent several limitations. The lRMSD is inherently hard to interpret, as medium values may stem from a fuzzy structural conservation contributed by all atoms, or from small regions that underwent large conformational changes while their complement is isometrically conserved. Possibly worse, the lRMSD involves the covariance matrix of centered atomic positions (see below), so that points far from the center of mass get more weight. This fact also has another consequence: by a packing argument, large proteins distribute atoms farther from the center of mass, so that the lRMSD depends on protein size. In order to weigh all points evenly and also to obtain a normalized measure, a variation of the RMSD obtained by restricting the calculation to unit vectors along the backbone and performing the optimization over rotations was developed [7]. Normalized alternatives were also proposed, respectively based on the radii of gyration of the molecules compared [8], and on normalization factors inferred from typical distributions of RMSD values [9, 10]. In a complementary line of attack, various superposition-free measures were proposed. To compare structures and models, one may use the lDDT which is an average value (computed over four thresholds) of fraction of distances within the chosen threshold [11]. Similarly, the contact area difference quantifies differences of contact areas between a model and a structure [12]. Finally, a normalized measure based on the Binet-Cauchy kernel, which inherently computes a scalar product between vectors defining the volume of tetrahedra was recently proposed [13].

Improvements were also reported to speed-up calculations. Efficient calculations targeting subsets of the aligned structures were investigated [14]. As an alternative to lRMSD calculations based on matrix decompositions–such as SVD, a fast determination of the optimal rotation matrix based on a Newton-Raphson quaternion-based method was reported [4]. Finally, recent work addressed the calculation of RMSD between flexible structures, the flexibility being modeled using collective coordinates obtained via normal mode analysis or PCA [15]. This latter work is especially interesting as it targets deformable structures modeled by means of collective motions.

To conclude, we also note that RMSD and lRMSD calculations are tightly related to the calculation of structural alignments between two structures. The intrication between scores and alignment methods was first exploited in [16], which performs an iterative alignment, guided by two scores, namely GDT (the fraction of residues (largest set, not sequence contiguous) that fit under a distance cutoff), and LCS (the fraction of amino acids defining the longest contiguous segment fitting under a given RMSD cutoff (positions of molecules fixed)). Since then, a variety of methods were proposed to detect and score structural motifs [17], including `DALI` [18], `TM-align` [19], `Apurva` [20], `LGA` [16], `Kpax` [21], or our persistence based method [22].

### 1.2 Contribution

In the sequel, we propose an elementary formula combining lRMSD measures, each associated with its own optimal rigid motion, so as in particular to assess molecular flexibility. With respect to the afore-discussed limitations of the lRMSD, our so-called *combined RMSD* has the following advantages:

- Flexibility is characterized by local structural alignments between structural motifs of the compared molecules, rather than with a global parameterization of motion.
- Flexibility can be assessed at multiple scales, by parameterizing the number of rigid motions used–this number is equal to the number of motifs mapped to one-another.
- The dependency of protein size is alleviated.

## 2 Combining independent lRMSD measures

### 2.1 Structures, alignments, and motifs

Let *A* and *B* represent either two distinct structures or two conformations of the same molecule or complex. Assuming these are proteins, we denote their number of amino acids *n_A_* and *n_B_*, respectively. Our structural comparisons are based on the coordinates of *particles*. In comparing two conformations of the same molecule, the particles may refer to all atoms, all heavy atoms only, or *C_α_* atoms; for two distinct molecules, we assume the particles are *C_α_* atoms.

Our comparisons shall use variants of the lRMSD, whose calculation requires an alignment of the two structures to be compared. We first recall:

#### Definition. 1

*Consider two sequences of length nA and n_B_. An alignment of length N* ≤ min(*n_A_,n_B_*) *is defined by two sets of indices I* = (*i*_1_,*i*_2_,…,*i_N_*) *with* 1 ≤ *i*_1_ ≤ *i*_2_ < ⋯ < *i_N_* ≤ *n_A_ and J* = (*j*_1_,*j*_2_,…*j_N_*) *with* 1 ≤ *j*_1_ < *j*_2_ < … *j_N_* ≤ *n_B_. An* alignment *is specified by the perfect matching {(*i*_1_,*j**_1_),…, (*i_N_,j_N_*)}.

Once the two structures have been aligned, abusing notations, we may reduce them to the two ordered point sets *A* = {*a_i_*}_*i*=1,…,*N*_ and *B* = {*b_i_*}_*i*=1,…,*N*_.

#### Structural motifs

To *localize* structural comparisons, we assume that *structural motifs* have been identified. Practically, two types of motifs may be used: features of proteins (domains, SSE), or motifs yielded by local structural alignment methods–see Introduction. More formally, we define:

##### Definition. 2

*Consider two structures A and B. A* motif *is a pair of set of particles M*^(*A*)^ ⊂ *A and M*^(*B*)^ ⊂ *S_B_ of the same size, together with an alignment between them*.

The alignment allows computing lRMSD(*M*^(*A*)^,*M*^(*B*)^). The fact that motifs may overlap calls for the following processing.

#### Motif graph for overlapping motifs

When several motifs exist for two structures, an important question is to handle them coherently. Since motifs may overlap, we define (Fig. 1):

**Figure 1:**
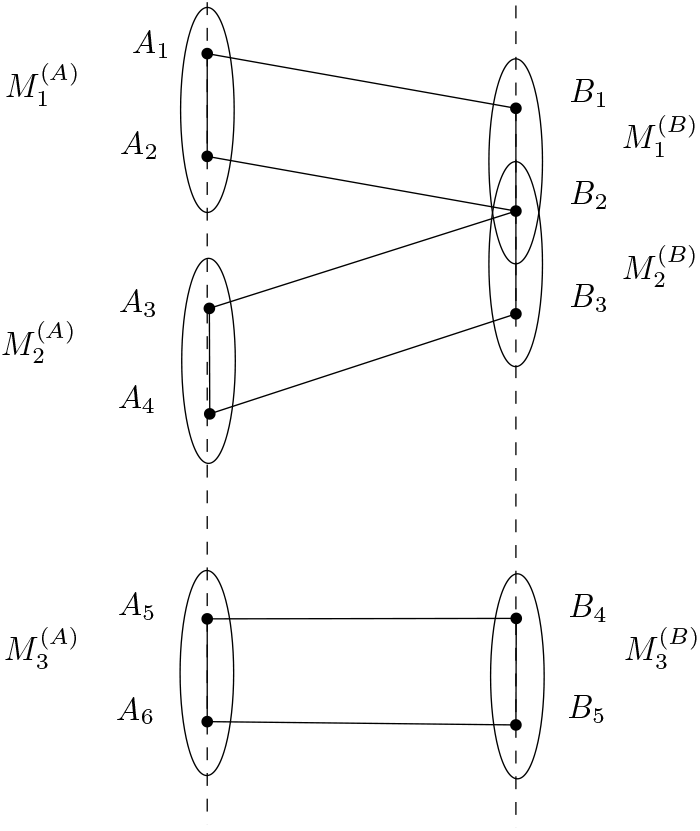
A motif graph (Def. 3). A toy system with two structures *A* and *B* involving 6 and 5 particles— say *C_α_*-s, respectively. There are three motifs, namely 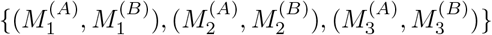. Motif edges are vertical edges connecting the particles; matching edges connect particles from the two structures. The three motifs induce two connected components, respectively containing 4 and 2 matching edges.

##### Definition. 3

*(Motif graph) The* motif graph *of a list of motifs* 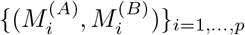 *is defined as follows: its node set is the union of the particles A and B; its edge set is the union of two types of edges*:

- matching edges: *the edges associated with the matchings defined by the motifs. NB: such edges are counted without multiplicity, that is, a matching edge present in several motifs is counted once*.
- motif edges: *edges defining a path connecting all amino acids in a motif*.

*Consider a connected component (c.c.) of the motif graph. Restricting each c.c. to each structure yields two subgraphs. The set of all such subgraphs is denoted* 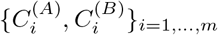.

The following observations can be made (Fig. 1). A subgraph (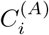 or 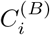) may not be connected. Also, the motif graph does not define, in general, a matching between the vertices associated with particles. More precisely, the *multiplicity* of a particle in a motif graph is defined as the number of edges incident to this particle. Despite these features, as we shall see below, the edges connecting the particles from 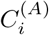 and 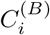 can be used to define a variant of the classical lRMSD.

### 2.2 Vertex weighted and edge weighted lRMSD

We introduce generalizations of the lRMSD and RMSD, using connected components of the motif graph. Before presenting these, we recall the construction of the weighted lRMSD (Def. 4).

#### Vertex weighted lRMSD : lRMSD_vw_

Consider two point sets *A* = {*a_i_*} and *B* = {*b_i_*} of size *N*. Also consider a set of positive weights {*w_i_*}_*i*=1,…,*N*_, meant to stress the importance of certain particles. The weighted RMSD reads as:

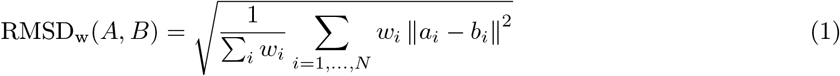

Let *g* a rigid motion from the the special Euclidean group *SE*(3). To perform a comparison of A and B oblivious to rigid motions, we use the so-called *least RMSD* [1]:

##### Definition. 4

*The* vertex weighted *lRMSD is defined by*

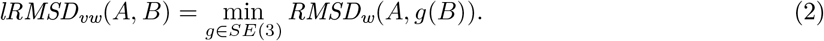

*The rigid motion yielding the minimum is denoted g^OPT^ (A,B) or g^OPT^ for short. The weight of the lRMSD_vw_ is defined as W_vw_*(*A, B*) = Σ*_i_ w_i_*.

Note that the celebrated lRMSD is the particular case of the previous with unit weights:

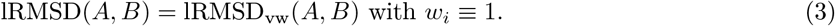

Denote R the sought rotation matrix [2] and *C* the covariance matrix

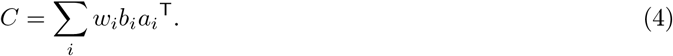

Upon centering the data, computing the lRMSD amounts to maximizing Trace(*RC*) [2, 4], a calculation which can be done with an SVD calculation.

#### Edge weighted lRMSD : lRMSD_ew_

Consider now the case where motifs have been defined for the two structures *A* and *B*. We wish to compare *A* and *B* exploiting the information yielded by the connected components of the motif graph (Def. 3). Consider the i-th c.c. of the motif graph. Let *e_i_* be the number of matching edges of this c.c. As usual, let *g*(*b_j_*) the position of atom *b_j_* from 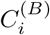 matched with atom *a_i_* from 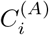, upon applying a rigid motion *g*. We define:

##### Definition. 5

*The* edge weighted lRMSD_ew_ *of the i-th c.c. of the motif graph is defined by*

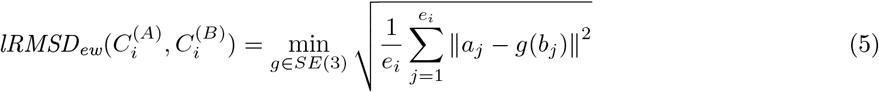

*The rigid motion yielding the minimum is denoted* 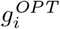.

*The weight of the lRMSD_ew_ is defined as* 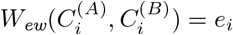.

To compute this quantity, we proceed as for the lRMSD_vw_, except that the covariance matrix from Eq. (4) is now obtained by summing over edges of the bipartite graph rather than on vertices. We also make the following

**Observation. 1** *If the motif graph defines a perfect matching between the particles of a connected component, then lRMSD_vw_ = lRMSD_ew_ for that component*.

### 2.3 Combined RMSD : RMSDcomb

Since the lRMSDew values are defined for each c.c. of the motif graph, we combine them to obtain a comparison of *A* and *B*.

Denote *m* the number of connected components of the motif graph and let *N_e_* = Σ_i_ *e_i_*. The edge weighted lRMSD can be combined into the following *edge-weighted combined RMSD*

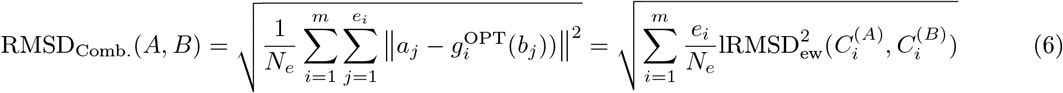

It is easily checked that the previous is a particular case of the following combined RMSD, which mixes individual lRMSD, be they vertex weighted (lRMSD_vw_) or edge weighted (lRMSD_ew_):

#### Definition. 6

*Consider two structures A and B for which non-overlapping regions* 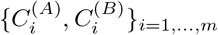 *have been identified – Def. 3. Assume that a lRMSD has been computed for each pair* 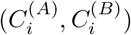. *Let w_i_ be the weights associated with an individual lRMSD. The* combined RMSD *is defined by*

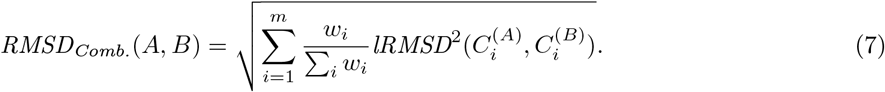

The following bounds are straightforwards convexity inequalities (proof in SI Sec. 7.1):

**Observation. 2** *The* combined RMSD *satisfies the following upper and lower bounds:*

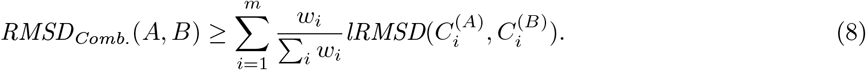

*Let* 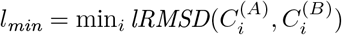 *and* 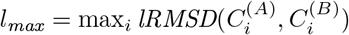. *One has*

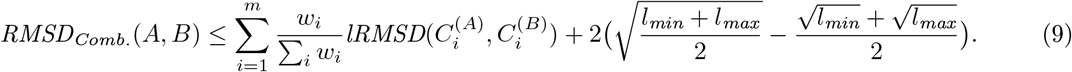

**Remark 1** *Equation (7) defines a RMSD rather than a lRMSD. To see why, observe that Eq. (2) defines a number which is the minimum of a quadratic optimization problem involving Eq. (4). Instead, Eq. (7) defines a number obtained by mixing solutions of such problems for connected components of the motif graph*.

**Remark 2** *Combined RMSD can be used iteratively. Consider two conformations of a a complex containing say two chains, each decomposed into motifs. In a first step, the combined RMSD exploiting the motifs can be used to compare the two instances of each chain across the two complexes. In a second steps, the combined RMSD of these combined RMSD can be computed. We illustrate this strategy to provide insights on novel conformations of quaternary complexes of hemoglobin in section 4.3*.

## 3 Implementation

All methods are available in the Structural Bioinformatics Library (http://sbl.inria.fr, [23]).

Three executables implementing the methods (sbl-rmsd-flexible-proteins.exe, sbl-rmsd-flexible-conformations.exe, sbl-rmsd-flexible-motifs.exe) are provided in the package Molecular_distances_flexible (https://sbl.inria.fr/doc/Molecular_distances_flexible-user-manual.html). Additional details are provided in SI Section 7.2.

Also of particular interest is the Structural_motifs package (see https://sbl.inria.fr/doc/Structural_motifs-user-manual.html), which makes available various methods to compute structural motifs [22].

## 4 Results

We illustrate insights yielded by the combined RMSD, which are out of reach for the classical lRMSD.

### 4.1 Assessing conformational changes: the example of a class II fusion protein

#### Biological context

Viruses replicate by hijacking the cellular apparatus of the host organism, with three main steps: entry into the host cell, multiplication, and egress of progeny virions from the host cell. For enveloped viruses, the entry requires the fusion of their envelope with the membrane of the host cell, a process triggered by a (low pH induced) conformational change of a membrane glycoprotein. Fusion proteins are ascribed to three classes denoted I, II and III, with class II fusion proteins typically structured three domains mainly composed of *β* strands. In their post-fusion conformation, these proteins act as trimers project out from the viral membrane. In the sequel, we use the combined RMSD to illustrate the conformation changes undergone by a prototypical class II fusion protein, from the tick-borne encephalitis virus. The ectodomain of this protein was crystallized both in soluble form (PDB: 1SVB, [24], 395 residues) and in postfusion conformation (PDB: 1URZ, [25], 400 residues) (Fig. 2). We use structural motifs computed using a method reported in a companion paper [22] to demonstrate the RMSD_Comb_. in such a setting (RMSD_MODE_MOTIF, SI Sec. 7.2.3).

**Figure 2:**
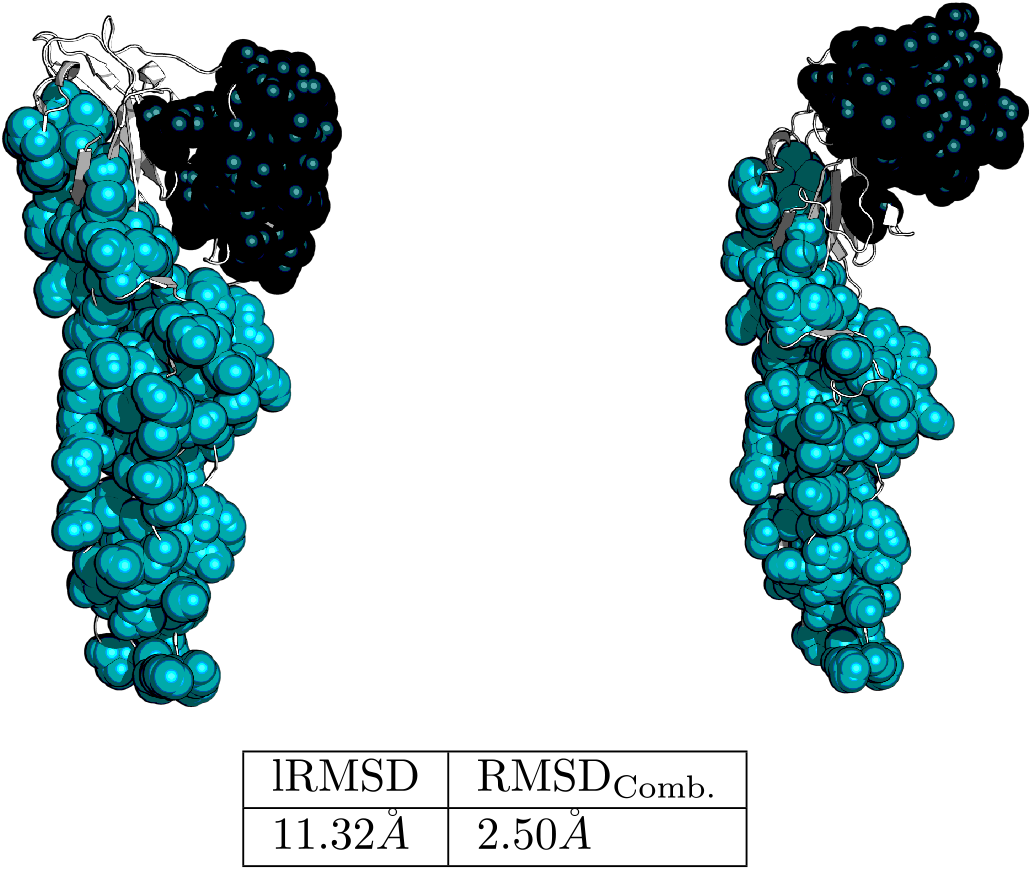
RMSD_Comb_. on overlapping structural motifs impervious to conformational changes: example on a class II fusion protein in soluble and post-fusion conformation. We display the two connected components, composed by the 31 structural motifs found by our method [22]. Most of the motifs overlap, which justifies a definition for overlapping motifs (Def. 3).

#### Results

At first glance, a global lRMSD calculation yields lRMSD = 11.32. To further assess the presence of rigid motifs moving relatively to one another, we computed motifs (Def. 2) using the method presented in the companion paper [22]. In a nutshell, this methods identifies structurally conserved motifs, each such region being a connected domain whose connectivity is ensured by pairs of atoms whose distance is conserved between the structures studied. (We note in passing that this strategy departs from the classical approaches targeting quasi-isometric motifs via the calculation of cliques [26].)

For the tick-borne encephalitis virus, the method yields a total of *p* = 31 structural motifs distributed in m = 2 connected components (|C_1_| = 109, |C_2_| = 51; Fig. 2). Finally, in computing the combined RMSD between the two connected components (Def. 3), one obtains RMSD_Comb_. = 2.50.

As illustrated by this example, the ability to identify structurally conserved motifs and to combine their lRMSD makes a dramatic difference in the overall comparison of two conformations. This problem is well known e.g. for the analysis of molecular dynamics trajectories, where erroneous assignments of structures to the same clusters / meta-stable states may jeopardize free energy calculations [27].

### 4.2 Building a phylogeny from structural data: the example of class II fusion proteins

#### Biological context

Following on the previous example, we now illustrate the interest of combined RMSD to build phylogenies of class II fusion proteins. As a dataset, we use 6 monomers of class II fusion proteins in their post-fusion conformation (Fig. 3). In order to assess the merits of the usual lRMSD and that of the combined RMSD to build phylogenies via dendograms, we carry out the analysis at two levels, namely using whole domains and the 22 motifs defined by SSE (Fig. 3(b)). (This corresponds to the setting RMSD_MODE_SEQ for homologous proteins, see SI Section 7.2.2.)

**Figure 3:**
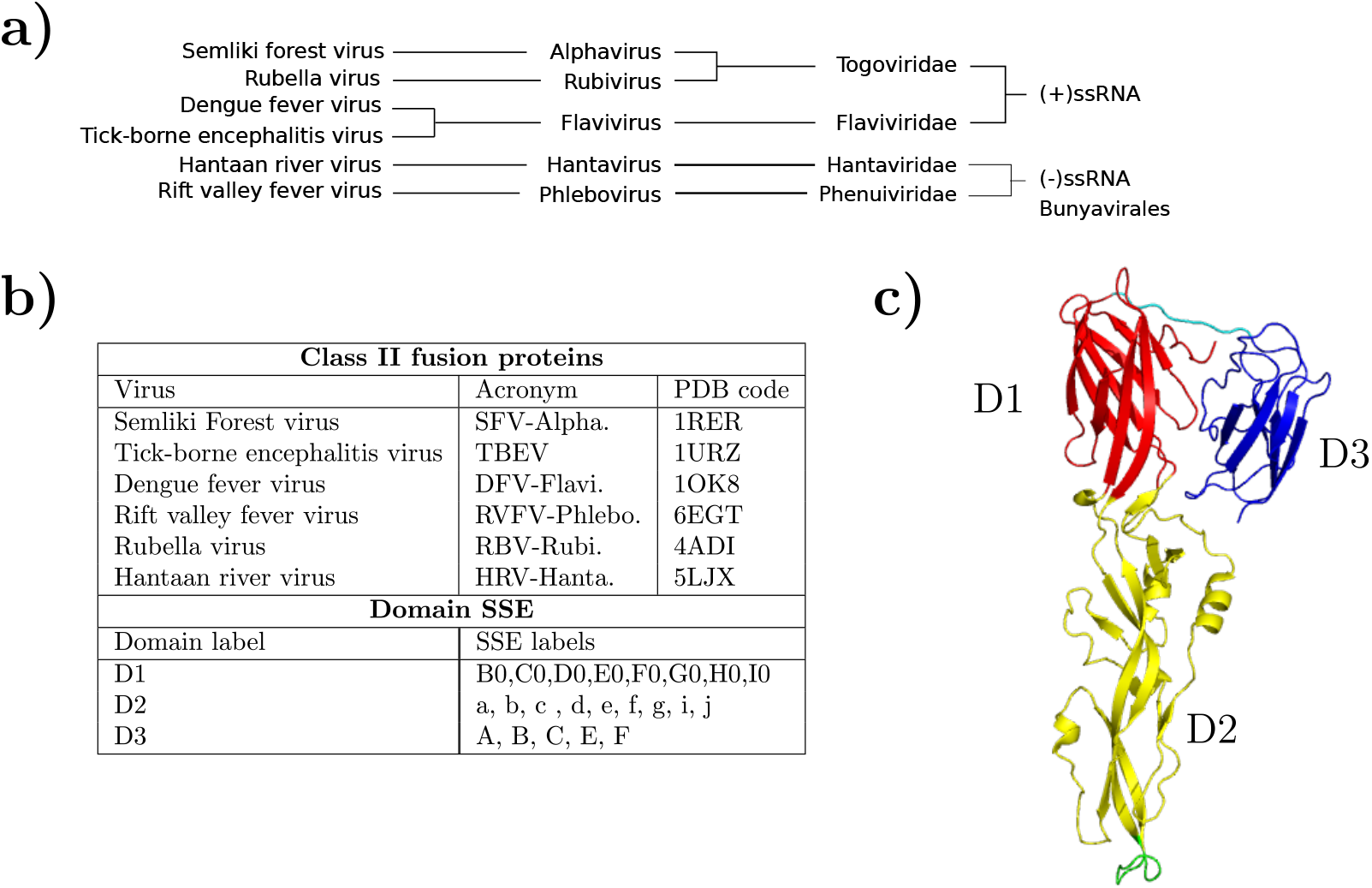
Class II fusion structures. **a**) Taxonomy of structures used in this study. **b**) Breakdown of the structures used in this study, as well as their domain and label labels. **c**) Domain decomposition of DFV-Flavi..

#### Results

Out of the six structures and for each pairwise comparison–15 of them, we used the executables provided in the `SBL` to compute: (1) a structural alignment using the `Apurva` algorithm [20]. This yields a lRMSD (Table 1), (2) the RMSD_Comb_. upon processing the regions defined by the three domains (Table 2), (3) the RMSD_Comb_. upon processing the regions defined by the SSE labels (Table 3). From the distance matrices displayed in the aforementioned tables, we perform three complete linkage hierarchical clusterings (Fig. 4). Each level of comparison conveys different information.

- **Full structure**. This level of information is enough to classify the two flaviviruses together. Out of the 6 structures described in Fig. 3, we know that 2 (HRV-Hanta. and RBV-Rubi.) have a domain swap. This adds considerable noise to the clustering.
- Domains. Here the two flaviviruses as well as the two bunyaviruses are clustered together, regardless of the domain swap.
- SSE. The two flaviviruses as well as the two togoviruses are clustered together, regardless of the domain swap.

**Figure 4:**
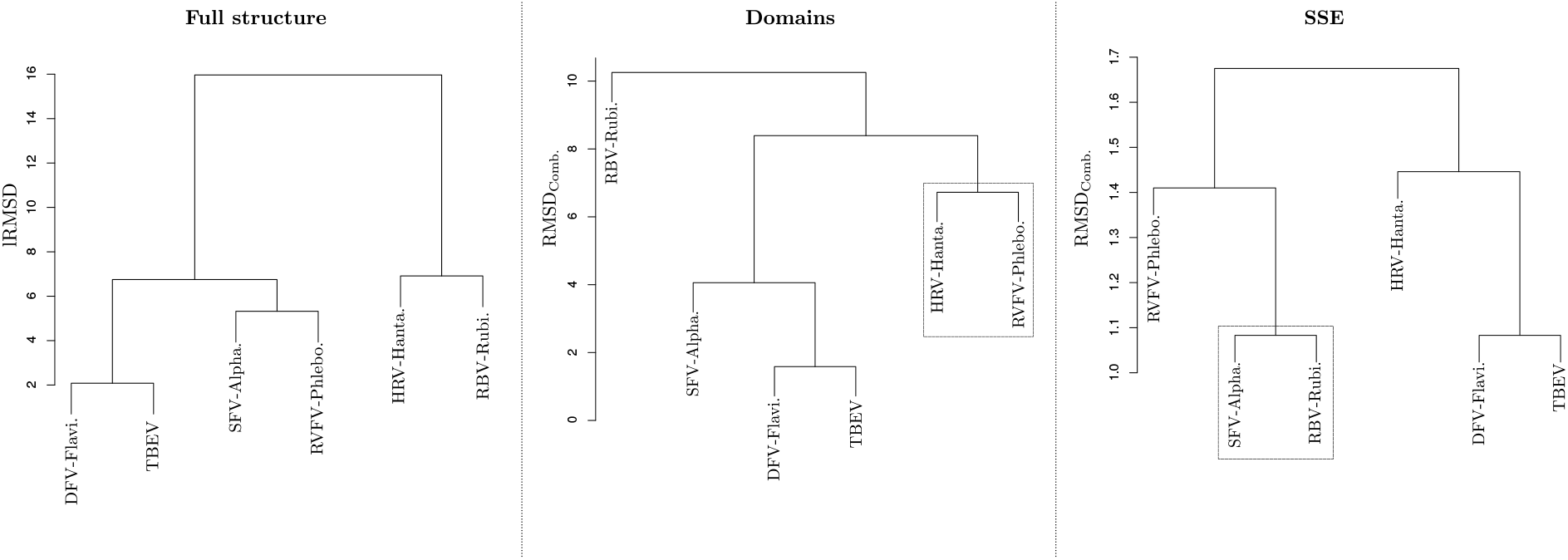
RMSD_Comb_. sharpens hierarchical clustering obtained for class II viral fusion proteins. Complete linkage hierarchical clustering of the structures defined in Fig. 3. (**Left**) Clustering obtained upon processing distances from Tab. 1. Global lRMSD after aligning structures with the Apurva algorithm. (**Center**) Clustering obtained upon processing distances from Tab. 2. RMSD_Comb_. using domains I, II and III. (**Right**) Clustering obtained upon processing distances from Tab. 3. RMSD_Comb_. using motifs corresponding to SSE.

To conclude, while the global lRMSD falls short from providing a satisfactory classification, combined RMSD fixes this limitation. One can further use the individual lRMSD value on a per domain basis to illustrate the structural diversity of these domains (SI Fig. 8).

### 4.3 Assigning quaternary structures: the example of hemoglobin

#### Biological context

Hemoglobin is the gas transporting metalloprotein in mammals. In humans, the predominant form of hemoglobin has a quaternary structure based on four subunits (SI Fig.9), namely two α chains (141 amino acids each), and two chains *β* (146 amino acids each). Each subunit contains a heme group consisting of a charged iron (Fe) ion held in an heterocyclic ring called porphyrin. It had long been believed that binding of O2 to one monomer triggered the transition of the tetramer from the tense (T) state to the relaxed (R) state, a mechanism at the core of cooperative binding [28]. Along this process, one pair of subunits rotates of an angle ~ 15deg about the other ([29] and SI Fig.9). However, the mechanism is more complex, and a third composite quaternary state, usually denoted R2 or Y was discovered long ago [30]. Based on these reference structures, a combination of various biophysical experiments [31] and macroscopic analysis (using angles and distances to qualify the quaternary structures) [32] had provided insights on gas binding by hemoglobin. Recently, these models were questioned by crystal structures which revealed that each tetramer captured hemoglobin in three quaternary conformations [33]. (NB: the three crystal structures are: half-liganded with phosphate (HL+, PDB: 4N7P), half-liganded without phosphate (HL−, PDB: 4N7O), and fully water-liganded met-hemoglobin with phosphate (FL+, PDB: 4N7N).)

Importantly, each new crystal was found to contain 3 new quaternary conformations denoted *A, B, C*. An assignment procedure based on difference distance matrices of the *α*_1_*β*_2_ subunits using R as a reference, (SI Sect. 7.4) supported the following: the A, B, C states respectively assumes states R2, R, and T. Moreover, visual inspection of the data [33, Fig. 2] shows that upon alignments, the tetramers tagged A or B superimpose almost perfectly, while those tagged C exhibit less coherent SSE (in particular helices from the FL+ crystal.

#### Results

As recalled above, assignment of quaternary states was mainly done so far using four rigid-body parameters related to the twofold symmetry [32], or using difference distance matrices with R as a reference state [33]. We revisit this problem, performing a complete hierarchical clustering of dimers using two combined RMSD : a combined RMSD mixing the lRMSD of the two chains; a combined RMSD mixing two combined RMSD –one for the 7 helices of chain *α* and one for the 8 helices of chain *β*. We use these combined RMSD to perform a hierarchical clustering (single linkage) of all combinations of α and β chains. Of particular interest are the clusterings of *α*_1_*β*_1_ and *α*_1_*β*_2_ dimers (Fig. 5, SI Fig. 10, SI Fig. 11), which evidence three interesting facts.

**Figure 5:**
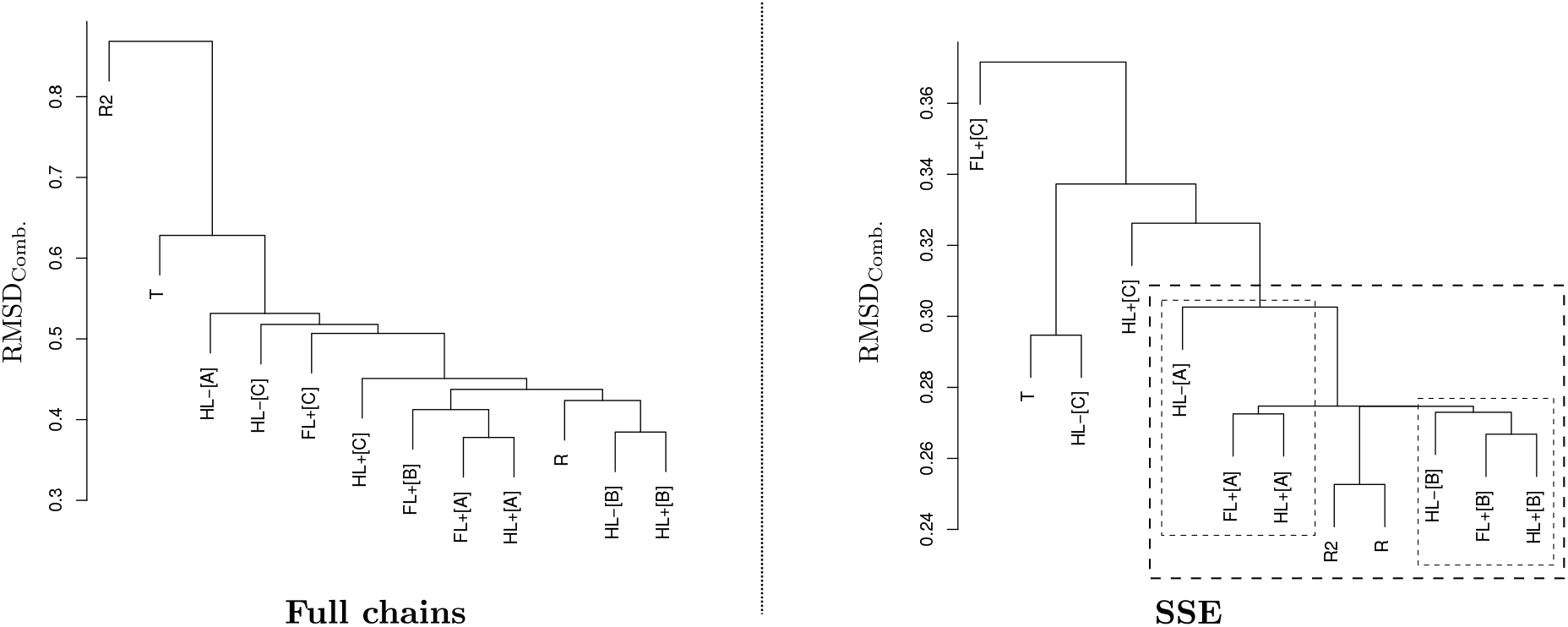
Assigning quaternary structures of hemoglobin using *α*_1_*β*_1_ dimers. The goal is to check which similarity measures allow one to cluster coherently the newly reported conformations *A, B, C* of hemoglobin tetramers ([33] and Sec. 4.3.), assumed to adopt quaternary structures corresponding to the R2, R and T states. The displayed hierarchical clusterings were built using the single linkage scheme. (**Left**) Using RMSD_Comb_. combining the lRMSD of the two chains *α*_1_ and *β*_1_. The hierarchical clustering obtained does not cluster coherently states A, B, C, and does not provide a coherent clustering with states R2, R and T either. (**Right**) Using RMSD_Comb_. combining the RMSD_Comb_. of the two chains *α*_1_ and *β*_1_, the former (resp. latter) based on the 7 (resp. 8) lRMSD between its helices. The clusters of conformations A, B and to a lesser extent C are well formed and coherent with the R2, R and T states.

**Figure 6:**
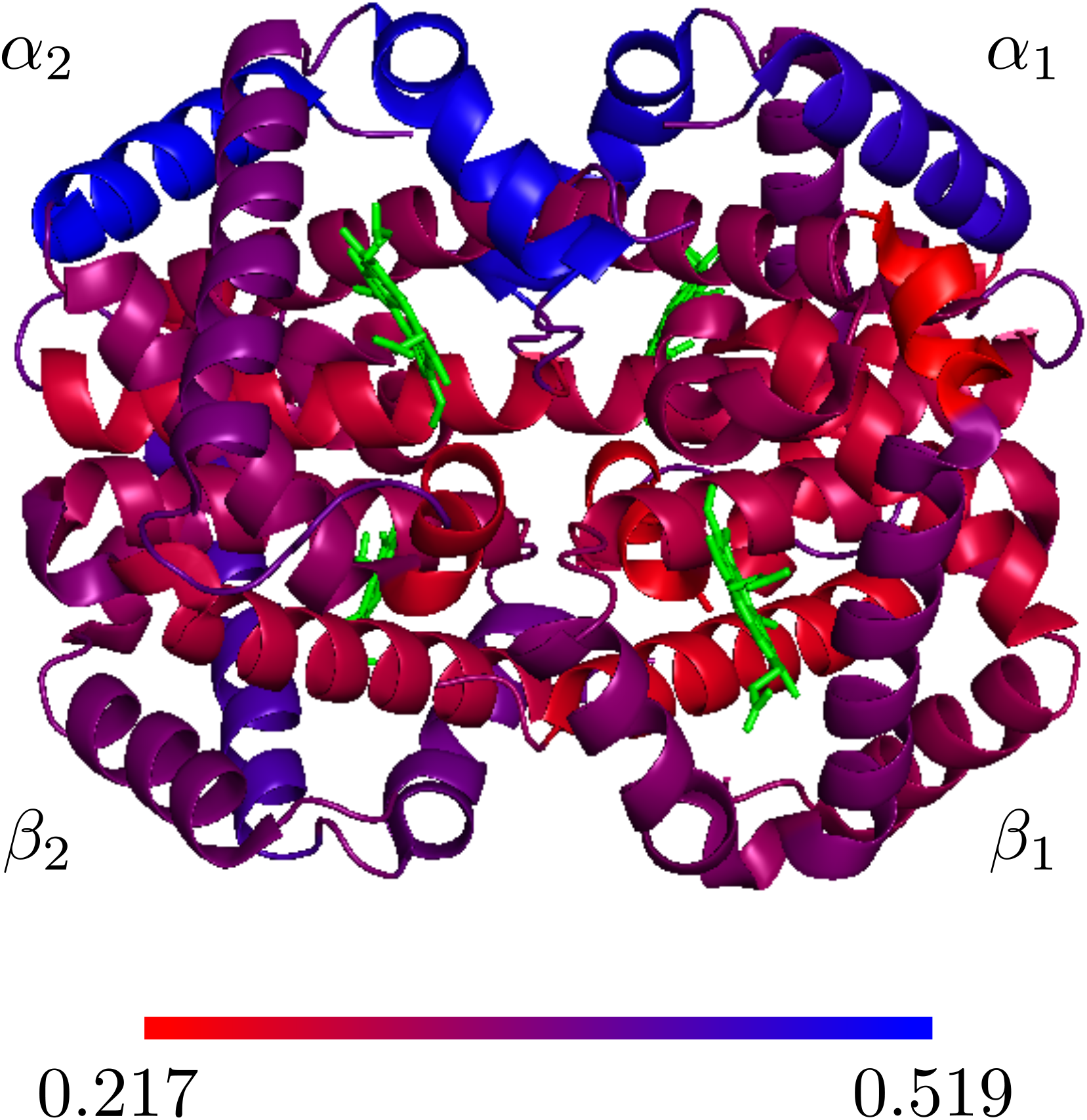
Structural conservation of hemoglobin. The *α* and *β* chains were respectively decomposed into 7 and 8 helices (Main text). For each helix, all pairwise lRMSD were computed using the 12 structures. Each helix was then color coded according to the gradient indicated. Visualization done with T conformation (pdbid: 2dn2).

First, let us consider the ability to group coherently the states *A, B, C*, versus states *R*2, *R* and *T*. The combined RMSD at the chain level is unable to separate the aforementioned conformations *A, B, C* (Fig. 5(Left), SI Fig. 10(Left)). On the other hand, the combined RMSD at the SSE level does retrieve the *A, B, C* conformations and groups them coherently with the *R*2, *R* and *T* states (Fig. 5(Right), SI Fig. 10(Right)).

Second, it confirms the information yielded by visual inspection, according to which conformations *C* of the tetramer are less coherent. Indeed, the hierarchical clustering isolates conformation C from the FL+ crystal (PDB: 4N7N). It also singles out HL− [A], which is less coherent with the other two [A] but also with *R* and *R*2 (SI Tables 4 and 5). This somewhat mitigates the analysis from [33], where HL− [A] is reported as a relaxed state between R and R2.

Third, the RMSD_Comb_. values required to form the clusters are tighter than those corresponding to RMSD values of clusters formed using the aforementioned angles and distances [32]. While a much larger dataset was used in this latter paper, the mean values reported for lRMSD between R2, R and T states are 0.40, 0.43 and 0.36, against worst case values ~ 0.27, 0.27 and 0.37 in our case.

Summarizing, the combined RMSD at the SSE level provides insights on the assignment of quaternary structures of hemoglobin, without using any reference state. Since SSE characterize quaternary structures, one can also assess the conformational changes undergone by helices in conjunction with the heme group (SI Fig. 6).

## 5 Discussion and outlook

The combined RMSD mixes independent lRMSD measures, each computed with its own rigid motion, therefore avoiding the global parameterization of conformational changes undergone by structures. Moreover, it can be computed at multiple scales, namely to compare secondary or tertiary structures based on structural motifs defined from the sequence or local structural alignments, or to compare quaternary structures from the same motifs. Finally, combined RMSD can be cascaded, so as for example to compare quaternary structures based on motifs defined from SSE elements. The notion of scale is in fact central to combined RMSD : on the one hand, the lRMSD computed for a whole structure typically yields an average signal shadowing the phenomenon scrutinized; on the other hand, computing a combined RMSD for too small structural elements would yield small values void of significance.

We illustrated the interplay between combined RMSD and pertinent scales on three non trivial examples, namely the analysis of large conformation changes, the design of phylogenies based on structural comparisons, and the identification of the quaternary structures of hemoglobin.

These examples may be discussed in the context of dynamical analysis of molecular machines, using the concepts of negative and positive discrimination of degrees of freedom. To articulate these notions, recall that two cornerstones of molecular simulations are move sets and collective coordinates. Move sets, on the one hand, are used in Monte Carlo methods and variants to generate conformations which are diverse and low in energy. Collective coordinates, on the other hand, are key to explore transition paths (and discover transient conformations), and compute free energy landscapes. Importantly, both concepts require understanding which degrees of freedom (dof) are key to account for the conformational changes studied.

To bridge the gap between move sets, collective coordinates, structural motifs, and combined RMSD, let us reconsider the scrutinized structures.

One the one hand, the example of the class II fusion protein undergoing a conformational change illustrates the notion of negative discrimination of dof. Indeed, the combined RMSD of the identified motifs being extremely small while the global lRMSD is large shows that in studying conformational changes between the two conformations studied, one can focus on those dof of atoms outside the motifs. In a sense, the combined RMSD rules out the dof of the motifs it qualifies.

On the other hand, the ability to cluster and qualify the quaternary structures of hemoglobin illustrates the notion of positive discrimination of dof. Indeed, while the global lRMSD does not yield any information, the restriction of lRMSD calculation to SSE, and the combination of the obtained values, yields valid biophysical classification. In other words, the combined RMSD positively discriminates the dof of the motifs it qualifies, calling for further studies to unveil the mechanism scrutinized. Such studies may focus on static analysis of crystal structures (detection of biophysical commonalities, formation/destruction of salt bridges, helix-to-coil transitions, etc). But they may also be dynamic, as the dof identified by may be targeted by complex move sets boosting the identification of structural intermediates.

Our methods to compute structural motifs and combined RMSD are made available within the Structural Bioinformatics Library (http://sbl.inria.fr). We anticipate that these tools will prove pivotal to conduct a wide array of structural analysis, both on static and dynamic structure of macro-molecules and their complexes.

## Abbreviations

RMSD: : root mean square deviation;
lRMSD: : least RMSD;
c.c.: : connected component.

